# Superimposed inhibitory surrounds underlying Saliency-based Stimulus Selection in avian Midbrain isthmi pars magnocellularis

**DOI:** 10.1101/2024.10.19.619184

**Authors:** Longlong Qian, Chongchong Jia, Jiangtao Wang, Shi Li, Songwei Wang

## Abstract

In the avian midbrain network, bottom-up spatial attention is directed by saliency-based stimulus selection. However, it remains unclear whether the isthmi pars magnocellularis (Imc), the first site in the midbrain network to represent stimulus selection, can represent stimulus salience and what is the mechanism by which the midbrain network computes salience. Here, we used two separate translational motion stimuli as the main stimulation protocols and conducted in vivo electrophysiological experiments in the pigeon’s Imc. By combining bio-plausible model validation, we found two types of inhibitory surrounds of the Imc neuron receptive field, homogenous inhibitory surrounds (HIS) and non-homogenous inhibitory surrounds (non-HIS), and elucidated the mechanism by which both arise. While HIS is local and depends on stimulus feature similarity, which can be used to compute stimulus saliency, non-HIS is global and doesn’t depend on stimulus feature similarity, which can be used to compute stimulus selection. Moreover, the superimposition of HIS and non-HIS modulates the neural response of Imc. The two inhibitory surrounds of Imc identified in this study more clearly elucidate the full process of achieving bottom-up stimulus selection based on saliency in the midbrain network and show that Imc is a nucleus that can represent both stimulus saliency and stimulus selection.

## Introduction

Finding the most salient targets in complex environments is important for survival activities such as feeding, courtship, and avoiding predators. Existing research suggests that the forebrain and midbrain (Mysore & Knudsen 2013; Sawant et al., 2022) are critical in stimulus selection (Bisley & Goldberg 2010; Squire et al., 2013; Asadollahi & Knudsen 2016; Mahajan & Mysore 2022). Among the midbrain spatial attention network focuses only on the relative priority of the stimulus and directs spatial attention to the highest priority location based on the saliency of the stimulus (Knudsen 2018; Schryver & Mysore 2019). However, how exactly the avian brain determines saliency remains uncertain.

The avian midbrain network consists of the optic tectum (OT, also termed the superior colliculus, in mammals) and nucleus isthmi, the latter including the nucleus isthmi pars magnocellularis (Imc), pars parvocellularis (Ipc), and pars semilunaris (SLu). The Imc receives excitatory projections from shepherd’s crook neurons (Shc, located in the OT 10 layer, OT10) and performs anti-topology inhibitory projection to the intermediate and deep layers of the OT (OTid) and other Imc neurons (Garrido-Charad et al., 2018; Wang et al., 2006; Wang et al., 2004). A cross-modal and global competition occurs in the OTid, a structure known to be involved in gaze control and attention (Mysore et al., 2010; Mysore & Knudsen 2011; Mysore et al., 2011; Mysore & Knudsen 2012). The selection of the OTid for the most salient stimuli relies on global inhibitory projections derived from Imc neurons (Mahajan & Mysore 2022; Mysore & Knudsen 2012). Schryver’s research showed that Imc neurons signal the strongest stimulus more categorically (Schryver et al., 2020), and earlier than the OT neurons. Mysore further demonstrated that the mutual inhibition between Imc neurons is an important neural mechanism for achieving stimulus selection (Mysore & Knudsen 2012; Schryver & Mysore 2023).

The saliency of the stimulus can be determined by how different it is from its history (that is, habituation, related to temporal saliency), how different it is from the surrounding objects (related to spatial saliency), and how relevant it is for the task at hand (Yoram 2015), where the first and second ones are bottom-up saliencies and the third one is top-down saliency. In this research, we focus on bottom-up saliency. According to Itti and Li’s research on saliency computation, the mechanism of saliency was considered to involve mutual inhibition between homogeneous feature detection neurons in the local region (no inhibition between non-homogeneous feature detection neurons) (Itti et al., 1998; Li 2016). Most researchers believe that the OT encodes stimulus saliency (Yoram 2015; Li 2016), and since this is one of the main factors influencing stimulus selection in the avian midbrain, Shc, the sole input to the nucleus isthmi at the OT (Garrido-Charad et al., 2018; Schryver & Mysore 2023), which integrates multilayered retinal ganglion cells (RGC) inputs originating from the OTs, can be considered to represent stimulus saliency. However, it remains to be determined whether Imc, the primary target nucleus for Shc projection to nucleus isthmi, also represents stimulus saliency, and what are the differences between the mechanism of stimulus saliency computation and that of stimulus selection.

Looming stimulus has been widely used to study stimulus selection in the midbrain, as it can effectively activate the neural response of each nucleus of the midbrain network (Asadollahi & Knudsen 2016; Mahajan & Mysore 2022; Schryver & Mysore 2019; Mysore et al., 2010). However, saliency must be expressed in terms of differences in interstimulus features, such as differences in motion direction, differences in stimulus flicker polarity, etc. The Looming stimulus has only intensity (velocity) information. So it is not possible to conduct research on saliency computation. The results of our previous studies (Wang et al. 2023) showed that Imc neurons are not only sensitive to translational motion targets but produce stimulus-specific adaptation (SSA) to translational motion targets with different directions (e.g., after repeated dorsal-ventral motion, ventral-dorsal motion is novel, and vice versa). This implies that stimuli moving in different directions are different features for Imc. Therefore, motion direction features can be used to conduct saliency-related studies.

Based on the above analysis, we used array electrodes to record the neural response of the Imc unit and conducted a study of Imc in pigeons using artificially generated visual stimuli. In the size-tuned stimulus model, the Imc response increases and then decreases with increasing stimulus size, and the inflection point of the response corresponds to a target size smaller than the range of the Imc excitatory receptive field, and we consider that the Imc neuron has an inhibitory surround overlapping with its excitatory receptive field. Using two separate translational motion stimuli, we found two types of inhibitory surrounds of Imc neurons, homogenous inhibitory surrounds (HIS) and non-homogenous inhibitory surrounds (non-HIS). While HIS depends on the similarity of stimulus features and is local in its scope of action in modulating Imc responses (The main range of action overlaps with the excitatory receptive fields of Imc neurons), it is thought to be computed on inhibitory projections between detectors of similar features upstream of Imc neurons and to play a role in the computation of stimulus salience in midbrain networks. Non-HIS does not depend on the similarity of stimulus features in modulating the responses of Imc neurons, is global in its scope of action, is thought to be computed on mutually inhibitory projections between Imc neurons and is used to compute stimulus selection in the midbrain network. Finally, we constructed a hierarchical coding model of RGC-Shc-Imc to verify the above hypothesis.

## Materials and methods

### Animal preparation

Given that sex differences in animals were rarely considered in researches that were conducted at the level of subcortical neurons and that most of the relevant works on midbrain network and Imc (Mysore & Knudsen 2013; Sawant et al., 2022; Mahajan & Mysore 2022; Schryver & Mysore 2019; Garrido-Charad et al., 2018; Mysore et al., 2010) do not pay attention to animal sex, the neuronal recordings in this study were conducted in male and female pigeons (Columba livia, 350–450-g body weight). The pigeons were housed in individual wire mesh cages under a 12:12-h light–dark cycle with free access to water and food. All experiments were conducted in accordance with the Animals Act, 2006 (China) for the care and use of laboratory animals and were approved by the Life Science Ethical Review Committee of Zhengzhou University.

### Surgery and recording

Experiments were performed following protocols that have been described previously (Wang et al., 2022; Wang et al. 2023). Briefly, surgery was initiated after the animals had been anesthetized with 20% urethane (1mL/100g body weight). Most existing experiments on the inhibitory surrounds of Imc and OT were performed under anesthesia (Schryver & Mysore 2023; Niu et al., 2022), therefore, experiments in this study were performed in anesthetized animals. When the birds closed their eyes and no longer responded to auditory or painful stimulation (mechanical stimulation such as pinching foot), meaning that they were in a state of deep anesthesia, they were transferred to a stereotaxic device. Their heads were placed in the stereotaxic holder, the right eye was kept open and moisturized with saline solution during the experiment, while the left eye was covered. A small hole was drilled in the bone to expose the left dorsal brain, allowing access to the Imc. A small slit was then made in the dura using a syringe needle, permitting the dorsoventral penetration through the Imc. Throughout the experiment, a heating panel was used to maintain the pigeon’s body temperature at approximately 41°C.

The multi-unit activity was recorded under urethane anesthesia using 16-channel microelectrode arrays (4 ×4, Clunbury Scientific, Bloomfield Hills, Michigan, USA), which were inserted into the Imc using a micromanipulator. We targeted the Imc following the previously described methods. Briefly, dorsoventral penetrations through the Imc were made at a medial-leading angle of five degrees from the vertical to avoid a major blood vessel in the path to the Imc. Identification of the recording site was based on the stereotaxic coordinates and expected physiological properties of Imc neurons: high firing rates and special visual RFs (to the best of our knowledge, Imc neurons are the only neurons in the midbrain that have a vertical elongated receptive field structure), and Imc targeting was finally validated at the outset of this study through anatomical lesions as described previously (Schryver & Mysore 2023; Wang & Frost 1991).

The spike signals of the units were recorded with a sampling frequency of 30 kHz and extracted with a bandpass filter (250–5 kHz). Local field potential signals were amplified (4000×), filtered (0–250 Hz), and continuously sampled at 2 kHz using a Cerebus® recording system (Blackrock Microsystems, Salt Lake City, UT, USA). All data recorded was analyzed off-line using MATLAB codes.

### Visual stimuli

Visual stimuli were generated using the MATLAB-based Psychophysics Toolbox (Psychtoolbox-3; www.psychtoolbox.org), running on Windows, and were synchronized with the recording system. An LED monitor (112 degrees vertical × 80 degrees horizontal, running at 100 Hz, PHILIPS, 558M1R) was placed tangentially to, and 40 cm from, the pigeon’s right eye to present monocular visual stimuli. The luminance of the gray screen background was 118cd/m^2^ and that of the black stimuli was 0.2 cd/m^2^.

Three visual stimuli protocols were used for data collection. The first was a black square (0.9 degrees) against a gray background, moving randomly at 90 degrees/s along a series of parallel paths (time window for the vertical and horizontal paths: 1.24 and 0.89 s, respectively) over the tangent LED monitor in front of the pigeon to map the RF of Imc units. Before the formal experiments started, the RFs of Imc units were estimated by manually sliding a dot delivered by a simple program around the screen while simultaneously monitoring the firing responses. Considering the large RF of Imc neurons, and to maximize the use of the monitor area, we then adjusted the position and angle of the monitor to ensure that its long axis was parallel with the long axis of the RF of the Imc.

The second visual stimulus, a black square ventral-dorsal moved at 30 degrees/s against a gray background, spaced by 15 degrees (the stimulus duration was 0.5s), was used to measure the size tuning curves of Imc units, and their sizes were 0.09, 0.15, 0.21, 0.27, 0.33, 0.39, 0.45, 0.6, 1.5, 3, 6, or 15 degrees. Within each trial, all the sizes were repeated five times following different pseudorandom orders, the intervals between each trial were 10s. All stimuli follow the same trajectory and pass through the center of the receptive field of the Imc units.

The third visual stimulus, two separated black squares (S1 and S2) of 3 degrees moved at 30 degrees/s against a gray background, spaced by 15 degrees (the stimulus duration was 0.5s), and their spatial distances were 3, 6, 9, 15 or 21 degrees. S1 was in the receptive field of the Imc units, and the position of S2 was on the left side of S1 (when the receptive field of the Imc unit was on the right side of the screen) or on the right (when the receptive field of the Imc unit was on the left side of the screen). There were two motion directions of the stimuli: dorsal-ventral and ventral-dorsal. S1 and S2 moved in the same or opposite direction at different spatial distances (20 cases). Within each trial, all the cases were repeated five times following different pseudorandom orders, the intervals between each trial were 10s.

### Histology

As described in our previous publications (Wang et al., 2022; Wang et al. 2023)), once data recording was complete, the actual site of the recorded unit in the brain was identified using anatomical lesions. Briefly, electrolytic lesions were made to mark the site of the recorded Imc neuron by passing a 0.2 mA current through the same recording electrode for 30 s. The bird was then euthanized with 20% urethane and perfused transcardially with phosphate-buffered saline followed by 4% paraformaldehyde. The brain was removed from the skull and tissues were postfixed for 12h, followed by immersion in 30% sucrose solution at 4°C for 24 h. Finally, the brain was frozen and cut into 50-µm transverse sections on a freezing microtome and stained with Cresyl violet for subsequent microscopic observations of marked and lesioned sites. The results show that all recorded units were in the Imc; therefore, no subject was excluded from the following data analysis.

### Analysis

The “Wave_clus” spike-sorting toolbox (Quiroga et al., 2004), a fast and unsupervised algorithm to the spike detection and sorting, was used to sort the recorded data into single units. We included in the analysis only those units that had less than 5% of the spikes within 1.5 ms of each other (Mahajan & Mysore 2018). The RFs of the Imc units were estimated by calculating the firing rate on the screen for each unit. The analysis window was from 80ms after the onset of the stimulus to 80ms after the offset of the stimulus, and the baseline response was estimated within 250 ms before the stimulus. Neural responses were quantified as the firing rate in the analysis window minus the firing rate of baseline activity and were calculated by averaging the response of the units across all trials; areas with responses exceeding the baseline were selected as the RFs of the recorded units.

The formulation of size tuning curve (Fig. 1e), Its parameters were calculated using SPSS, the equation is as follow:

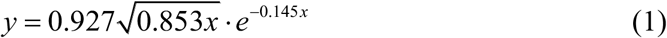

**Fig. 1.**
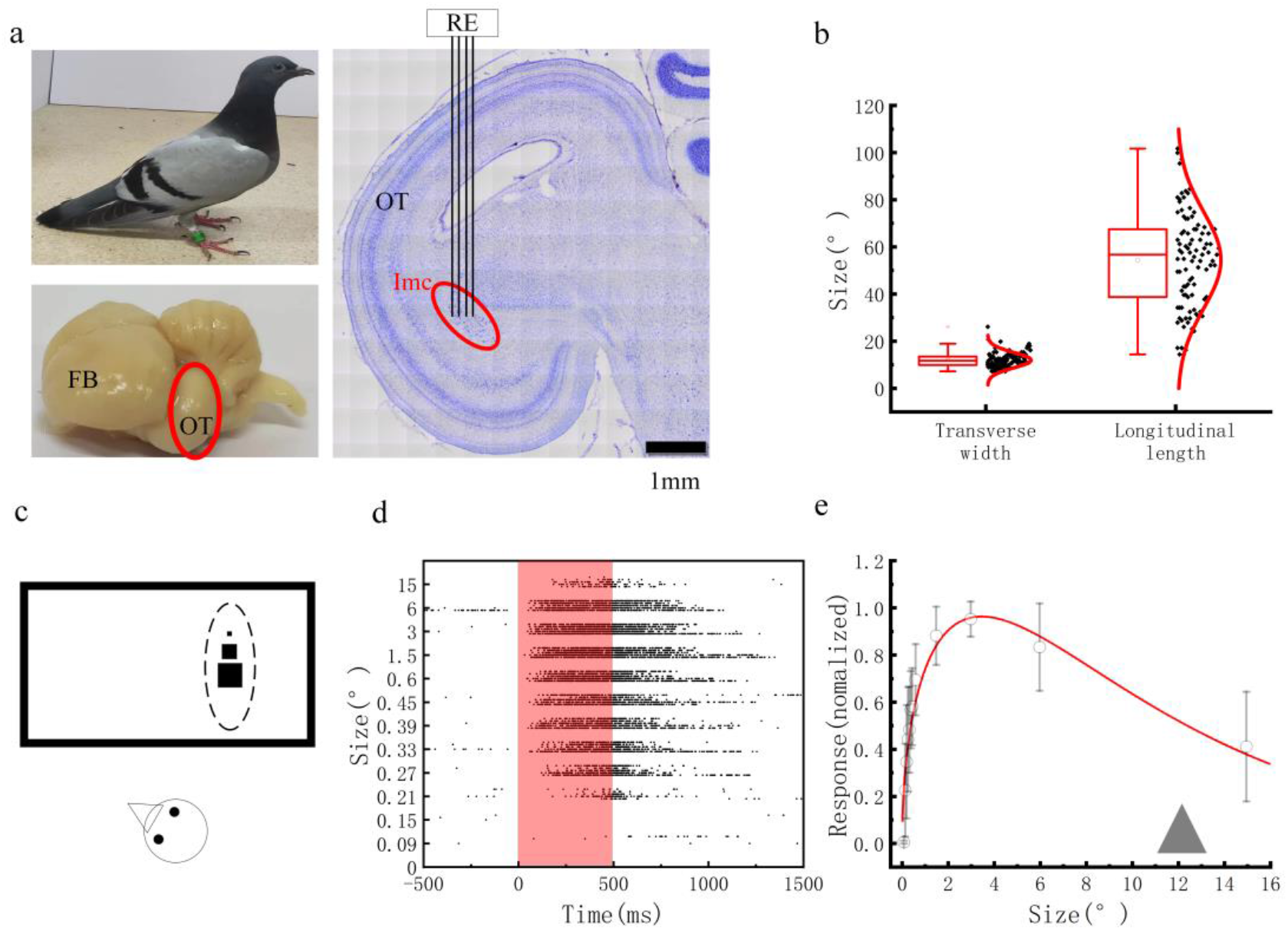
Receptive field properties and size tuning of the lmc. a. Top left, pigeons were used as experimental animals in this research. Top right, the Imc was encapsulated by the OT (the red elliptical area is OT). Right, electrodes were implanted from the upper OT, and the site of electrode implantation is shown in the red ellipse, which is the position of the Imc. b. The transverse width and longitudinal length of the Imc unit RF, where the transverse width of the Imc unit was obtained by the translational motion stimulus of dorsal-ventral and ventral-dorsal, the transverse width of the RF is 12.27°±3.59°(n=99), and the longitudinal length of the RF was measured by the translational motion stimulus of nasal-temporal and temporal-nasal (59.44°± 22.64°, n=99). c. Stimulus protocols of size-tune, S1 is inside the RF (S1 size = 0.09°, 0.15°, 0.21°, 0.27°, 0.33°, 0.39°, 0.45°, 0.6°, 1.5°, 3°, 6°, and 15°). d. The raster plot of an example Imc unit to S1 of different sizes; the shaded red ranges represent the stimulus duration. e. The normalized response of the Imc unit population to targets with different sizes; the red curve is a fitting of the Imc unit population average FR, (n=68, MSE = 0.0923); data represent ±SEM.

The influences of the Imc’s directional-HIS (blue curve, SD-OD) and OIS (red curve, OD-Single) on the Imc response under S1 and S2 at different distances were shown in Fig. 2h (the example Imc unit) and 2k (the Imc units population), these curves were fitted by using the Gaussian function in SPSS (the parameters are shown in Table 1). The equation is as follow:

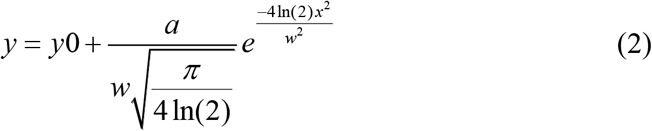

**Table 1.**
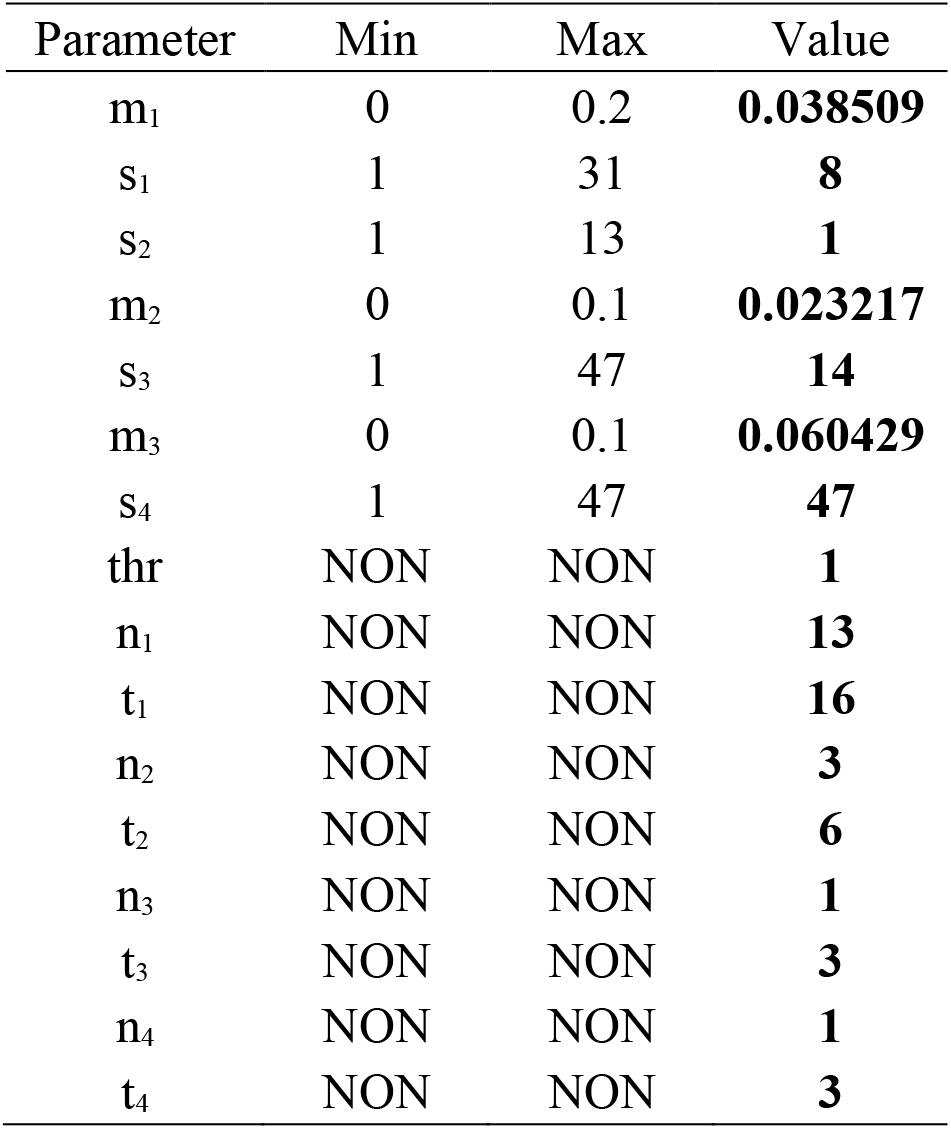
The parameter corresponding to the curve in Fig. 2h and 2k.

**Fig. 2.**
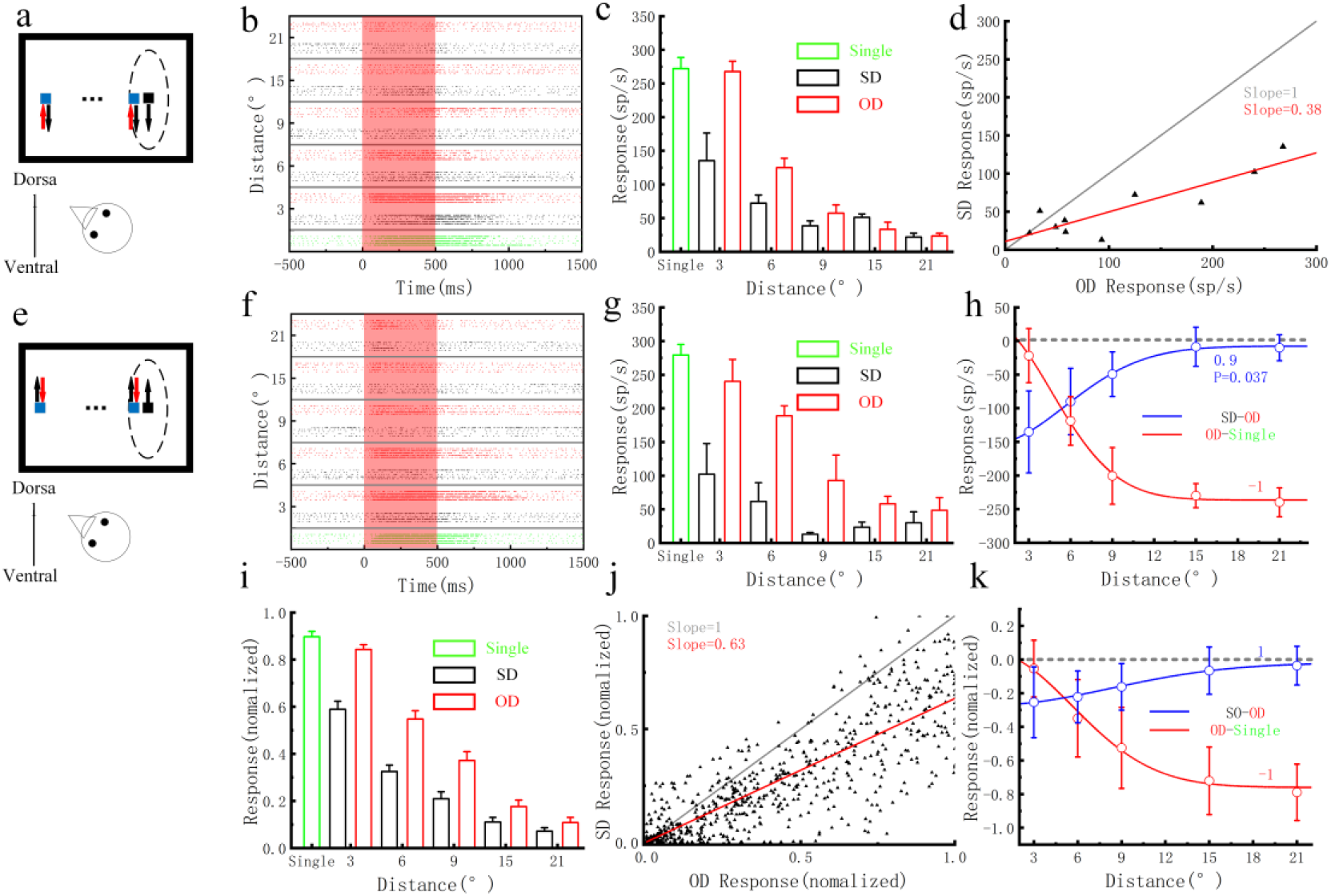
Directional homologous inhibitory surrounds and other inhibitory surrounds of the lmc. a and e. The stimulus protocols for studying the properties of the HIS (direction) and OIS of the lmc, in which the target size is 3°, the stimulus S1 (black) in the center of Imc units RF, the competitive stimulus (S2, blue) appears at 3°, 6°, 9°, 15°, and 21°away from S1, the motion directions of S2 are dorsal-ventral and ventral-dorsal, the directions of S1 are dorsal-ventral and ventral-dorsal in subplot a and e, respectively. b and f. The raster plots of the responses of an example Imc unit (corresponding to a and e, respectively), the data of S1 and S2 at different distances are separated by solid gray lines, where the green data are the response when single S1 (single), the black data indicate that S1 and S2 moving in the same direction (SD), red data indicate that they moving in the opposite direction (OD), and the red shaded range indicate the stimulus duration. c and g. The average FR of the example Imc unit during the stimulus duration in b and f, respectively. d. Scatter plot of the Imc unit response when S1 and S2 in the SD (ordinate) and OD (abscissa) conditions, the solid red line is the straight line fit of the data (slope = 0.38 ±0.074). h. The influence of the Imc’s directional-HIS (blue curve, SD-OD) and OIS (red curve, OD-Single) on the example Imc unit response under S1 and S2 at different distances. The curves are fitted by using the Gaussian curve, the correlation coefficients (response versus competitor size) are computated by Spearman test (red data is −1, n = 5; blue data is 0.9, P = 0.037, n = 5). i. Population summary (Imc), conventions as in c and g (n=67). j. Scatter plot of the response of the Imc unit population when S1 and S2 move in the SD (ordinate) and OD (abscissa) conditions, and the solid red line is the straight line fit of the data (slope = 0.63 ±0.016). k. Population summary (Imc), conventions as in h, the curves are fitted by using the Gaussian curve, the correlation coefficients (response versus competitor size) are computed by Spearman test (red data is −1, n = 5; blue data is 1, n = 5).

The Spearman test was used for the calculation of the correlation coefficient involved in this paper. The linear fit in Fig. 2 was achieved using the SPSS software.

## Model

### Hierarchical coding model of RGC-Shc-Imc

Next, to assess our hypothesis regarding the mechanism of the Imc’s directional-HIS and OIS, we constructed an Imc hierarchical neural computational model. To simplify the calculation, only the Imc’s HIS based on the characteristics of the direction of motion is constructed. The inhibitive projection of the Imc to Shc and between the Imc is used to construct the Imc’s non-HIS. The reliability of the model is verified using a video as input. The model consists of three layers: RGC, L10 (Shc), and Imc, were shown in Fig. 3.

**Fig. 3.**
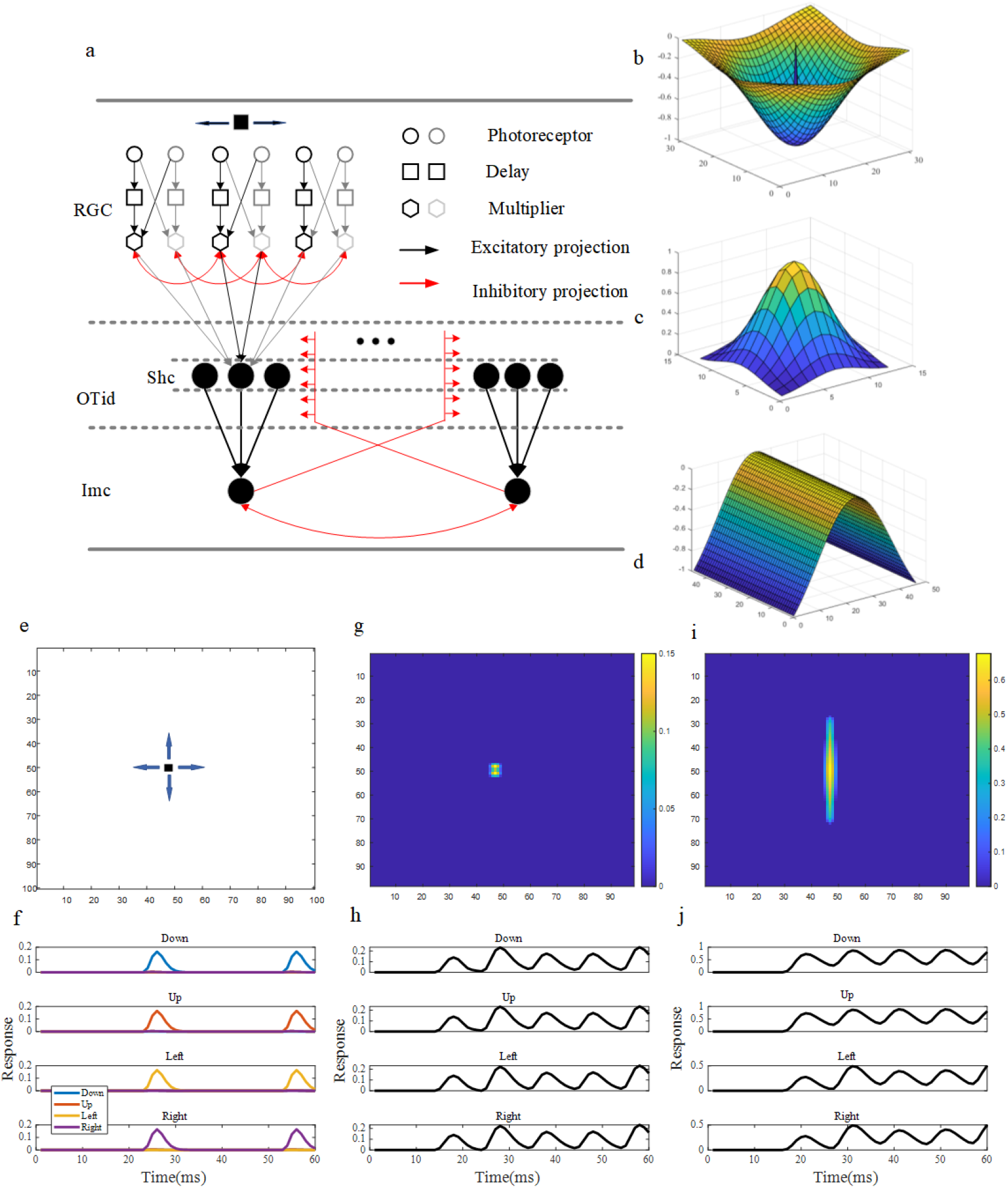
Imc’s hierarchical neural computational model. a. Diagram of Imc’s hierarchical neural computational model. b. The kernel function of the inhibitory projection between homologous RGCs. c. The kernel function of RGCs project to Shc. d. The kernel function of inhibitory projections of the Imc to Shc and between Imc neurons. e. The validation of the stimulus protocols for the model, in which the arrows indicate the direction of motion of the stimulus. f. The response of model RGC neurons to translational motion stimuli, in which the four subplots from top to bottom represent the neural responses of different types of motion detectors when the target is in downward, upward, leftward, and rightward motion, respectively. The blue, red, yellow, and purple curves represent the neural responses of the downward, upward, leftward, and rightward motion detectors, respectively. g. The response of model Shc neurons layer. h. The response of the model Shc, in which the four subplots from top to bottom represent the neural response when the target downward, upward, leftward, and rightward passes through the RF of model Shc neurons. i. The response of model Imc neurons layer. j. The response of model Imc neurons, in which the four subplots from top to bottom represent the neural response when the target downward, upward, leftward, and rightward passes through the RF of model Imc.

### Retinal layer

Because the stimulus protocols involved in this experiment were all translational motion stimuli, only motion detection neurons were constructed in the RGC layer. In this study, an elementary motion detector was used to realize motion detection in different directions (Reichardt 1961). The elementary motion detector was widely used to model the motion detection neurons of insects, and its basic principle was shown in Fig. 3a. First, the brightness change was detected through the photoreceptor (ON and OFF photoreceptor). When the ON photoreceptor detected an increase in brightness or when the OFF sensor detected a decrease in brightness, the photoreceptors generate a unit signal, that are *P* ^+^ and *P* ^−^.

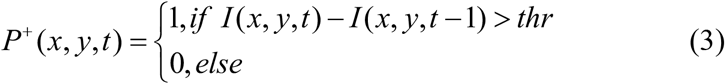

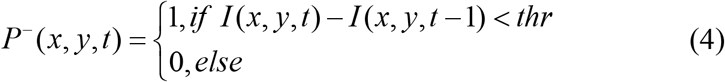

where *thr* is threshold of brightness change, (*x, y*) represents localtion of sensor, *t* is time, *I* is the input image. In this paper, motion detector neurons of the leftward(*R_L_*), rightward(*R_R_*), upward(*R_U_*) and downward(*R_D_*) were constituted.

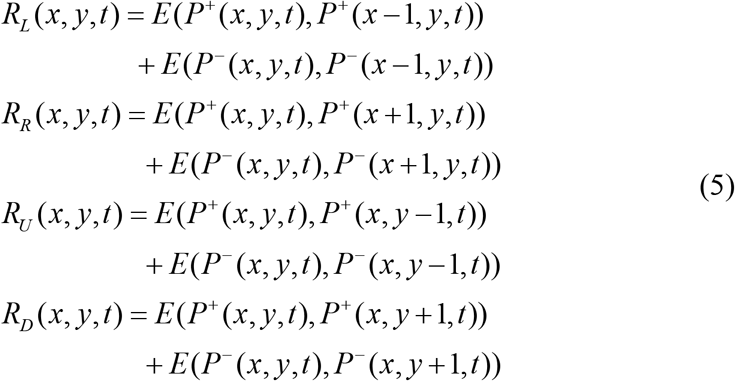

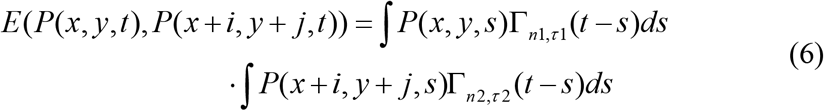

where *i, j* ∈ {−1,0,1}. Here the Gamma function Γ_*n,τ*_(*t*) was used to calculate the postsynaptic membrane potential after the detected brightness change reached the threshold (Wang et al., 2018).

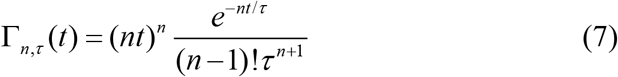

where *n* is orders of the Gamma kernels while *τ* is time constants.

### Shc layer

Existing studies have shown that Shc neurons receive input from the superficial OT layer (retinal output layer) (Wang et al., 2004; Garrido-Charad et al., 2018). Based on the previous analysis, Imc’s directional-HIS is computed in upstream of Shc, and to simplify the calculation, we directly model it as a mutual inhibition between homogeneous RGCs at the synapse of the RGC-Shc junction (using a Gaussian function to describe this inhibition). The Imc has an inhibitory projection of antitopology to the deep layers of the OT (Wang et al., 2004), and this antitopological projection is the key to achieving “winner-take-all” (Mahajan & Mysore 2022), which is modeled here as an inhibitory projection of the Imc to Shc.

First, the mutual inhibition between homogeneous RGC outputs was modeled as follows:

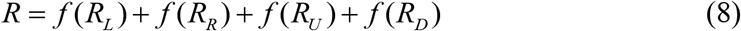

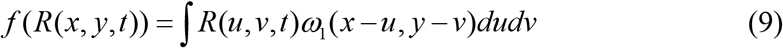

where *ω*_1_ = *µ*_1_*nom*(−*G*_1_) is the kernel function of mutual inhibition between homogeneous RGC outputs (shown in Fig. 3b). 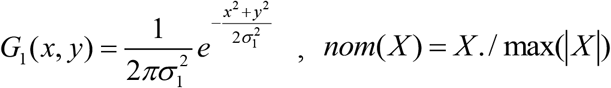, its function is to normalize kernel functions, the central position of *ω*_1_ is set to 1, that is, the RGCs are not inhibitory themself.

The response of the Shc was modeled as follows:

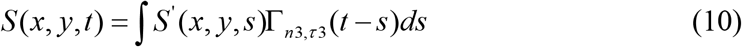

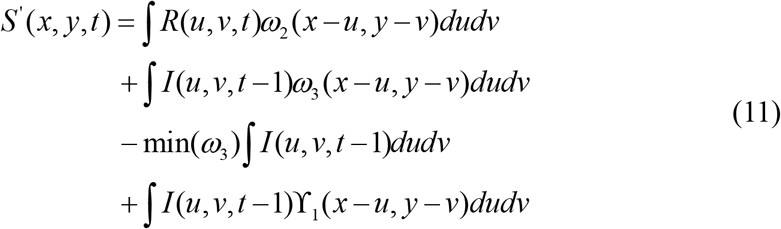

where *ω*_2_ = *nom*(*G*_2_) is the kernel function of RGC project to Shc (as shown in Fig. 3c), 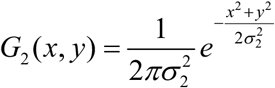. The modeling of the projection of Shc receiving Imc were divided into two parts, one is the region where the inhibitory projection weight gradient (the second term behind the medium sign in Equation 11), that is, *ω*_2_ = *µ nom*(*G*_3_) (as shown in Fig. 3d), 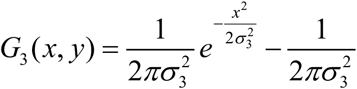. The other part is the region where the inhibitory weights are stable (the third and fourth terms behind the medium sign in Equation 11), that is, min(*ω*_3_) (the minimum value in matrix *ω*_3_), γ_1_ is an all-1 matrix of the same size as *ω*_3_.

### Imc layer

The Imc plays a key role in the winner-take-all process performed by the midbrain network, which directly determines the gaze position at the next moment. The Imc receives excitatory projections from Shc, and there are mutual inhibitory projections between the Imc (Mysore & Knudsen 2012). As the the excitatory receptive field of the Imc is vertical and elongated, to simplify the calculation process, an Imc neuron receives the excitatory projections of several Shc neurons in the vertical distribution corresponding to the receptive field space of the Imc neuron and constructs inhibitory projections like the Imc to Shc between Imc neurons. The membrane potential function of Imc neuron was modeled as follows:

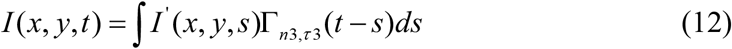

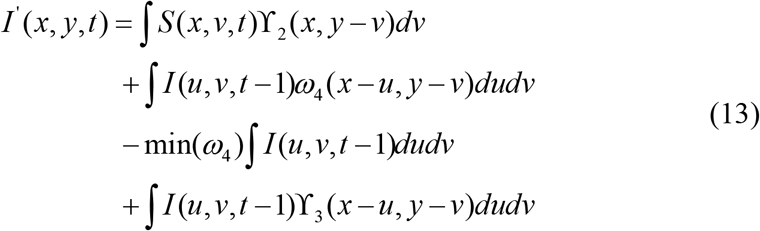

where γ_2_ is an all-1 matrix of *N* ×1, indicates that Imc receive input from N Shc, and the mutual inhibitory projections between Imc are divided into two parts, one is the region where the inhibitory projection weight gradient (the second term behind the medium sign in Equation 13), that is, *ω*_4_ = *µ*_3_*nom*(*G*_4_) (as shown in Fig. 3d), 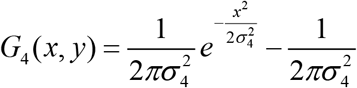. The other part is the region where the inhibitory weights are stable (the third and fourth terms behind the medium sign in Equation 11), that is, min(*ω*_4_) (the minimum value in matrix *ω*_4_), γ_3_ is an all-1 matrix of the same size as *ω*_4_.

### Parameter optimization

The differential evolution (DE) algorithm was used to optimize the parameters involved in the model. DE is an efficient and high-performing optimizer for realvalued parameters (Salt et al., 2019; Storn & Price 1997). As it is based on evolutionary computing, it performs well on multimodal, discontinuous optimization landscapes. Storn and Price (Storn & Price 1997) showed that their original DE outperformed several other stochastic optimization techniques in benchmarking tests while requiring the setting of only two parameters, crossover probability CR and differential weight F (there CR=0.7, F=0.5).

The initial population, *X*_1_ = {*x*_1,1_, *x*_2,1_,…, *x*_*N*,1_}, where N=20 is the size of the population, *x*_*i*,1_ ∈ℛ^*D*^ is an individual that contains the D=7 parameters to be optimized, is generated from random samples drawn from a uniform probability distribution of the parameter space, bounded to the range of the respective variable. These bounds are shown in Table 2. The fitness of each vector in the population is calculated by the objective function, the optimization function is formulated as:

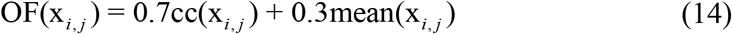

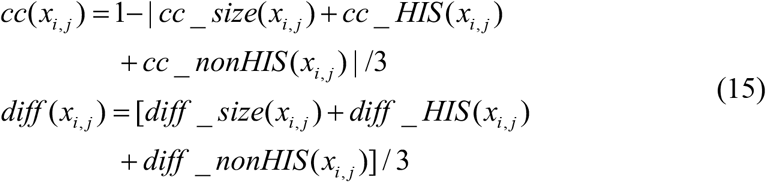

**Table 2.**
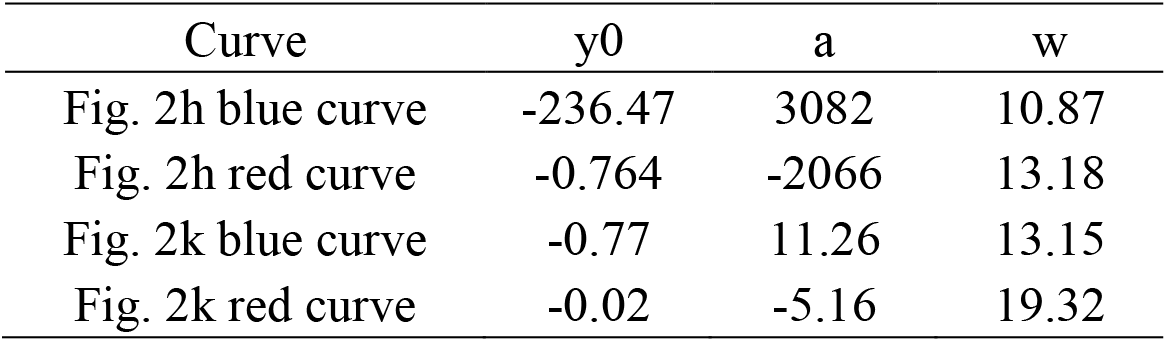
The range of parameter values and the optimization results involved in the Imc’s hierarchical neural computational model.

Where *cc* _ *size*(*x*_*i, j*_), *cc* _ *HIS* (*x*_*i, j*_) and *cc* _ *nonHIS* (*x*_*i, j*_) represent the correlation coefficient, *diff* _ *size*(*x*_*i, j*_), *diff* _ *HIS* (*x*_*i, j*_) and *diff* _ *nonHIS* (*x*_*i, j*_) represent mean response difference between the model Imc and Imc unit(regarding size tuning, Imc’s HIS, and Imc’s non-HIS, respectly). Evolution stops when evolution exceeds 100 generations, or the objective function is less than 0.01.

## Result

Data presented here were collected from single- and multi-Imc units in 9 anesthetized pigeons. The spike and local field potential signals of all units were stored simultaneously throughout the experiments. The multi-unit data were then subjected to spike sorting (see Materials and Methods) for all remaining analyses in this study.

### The properties of excitatory receptive field and size tuning of the Imc

In this study, we used pigeons as the experimental animals (Fig. 1a), an electrode array was implanted from the upper part of the OT (Fig. 1a). The recorded neurons were determined to be Imc units by the coordinate position of the stereotaxic apparatus and the morphology of the excitatory receptive field (ERF, the ERF of Imc neurons was long and narrow longitudinally distributed (Wang & Frost 1991; Wang et al., 2022; Wang et al., 2023)), and the implantation site was histologically identified after the experiment. When measuring the ERF of the Imc unit, the translational motion stimuli in four motion directions were used as stimulus protocols, with an average transverse width of approximately 12.27°± 3.59°(n=99) and a longitudinal length of approximately 59.44°± 22.64°(n=99) (as shown in Fig. 1b). According to our previous work (Wang et al., 2022), the Imc prefers dorsal-ventral and ventral-dorsal motion, so translational motion targets were used in this study.

To further investigate the sensitivity of the Imc to translational motion target size, a size-tuned stimulation protocol (Fig. 1c) was designed, and the target sizes (square) involved in the stimulation mode were 0.09°, 0.15°, 0.21°, 0.27°, 0.33°, 0.39°, 0.45°, 0.6°, 1.5°, 3°, 6°, and 15°. Fig. 1d shows the raster plotting of example Imc unit to targets with different sizes. Here, the average firing rate (FR) of the stimulus duration was used to measure the sensitivity of the Imc unit to targets with different sizes. The results of the normalized response of the Imc units population (n=68) to targets with different sizes were shown in Fig. 1e. The results of the average FR fitting indicated that the responses of the Imc units increased and then decreased with the increasing target size, and the target size (3°) corresponding to the inflection point of the response was smaller than the average transverse width of the Imc unit ERF (12.27°± 3.59°, n=99), indicating that there was an inhibitory surround that affects the response of Imc neurons within the range of the ERF of Imc.

### Imc’s directional homologous inhibitory surrounds and other inhibitory surrounds

According to Itti and Li’s work on saliency computation (Itti et al., 1998; Li 2016), mutual inhibition between similar feature detection neurons (homologous neurons) in a local area is the mechanism of saliency; that is, the neural response will be weakened when two stimuli with the same features are present simultaneously compared to when only one stimulus is present. On this basis, we ask whether the response of Imc neurons satisfies the above conditions, or whether Imc units represent saliency.

To this end, two stimulus protocols (Fig. 2a and 2e) were designed. In Fig. 2a, the stimulus S1 was located in the center of the ERF of Imc unit (the motion direction of S1 was dorsal-ventral), and the competing stimulus S2 was placed to the side of S1 (the motion direction of S2 was dorsal-ventral and ventral-dorsal). To avoid the influence of the motion direction of the target on the experimental results, the stimulation protocol shown in Fig. 2e was used as a supplementary experiment. The motion direction of S1 was ventral-dorsal, the motion directions of S2 were dorsal-ventral and ventral-dorsal, and the position of S2 was on the left side of S1 (the receptive field of the Imc unit is on the right side of the screen) or on the right (the receptive field of the Imc unit is on the left side of the screen). The parameters used to fit the curves to the results are shown in Table 1.

The responses of the example Imc unit to the above stimulus protocols were shown in Fig. 2b and 2f, and the average FR of the stimulus duration was used to measure the strength of response of the Imc units under different stimuli (Fig. 2c and 2g). To compare the differences in the responses of the example Imc unit between two stimuli (S1 and S2) moving in the same direction (SD) and in the opposite direction (OD), we plotted the scatter with the response in the opposite direction condition as the abscissa and the response in the same direction condition as the ordinate (Fig. 2d). The scatters were fitted in a straight line (slope = 0.38), and the results showed that the responses of the example Imc unit in the same direction condition were significantly weaker than those in the opposite direction condition. To test the generality of the experimental phenomenon, the response results of the recorded Imc unit population (n = 68) were normalized (as shown in Fig. 2i). The Imc units population responses in the same direction and the opposite direction condition were plotted as a scatter plot on the coordinate axes in Fig. 2j, and a linear fit was applied to all of the data (slope = 0.63). The results showed a widespread occurrence in which the responses of Imc units in the same direction condition were weaker than those in the opposite direction condition. This phenomenon exhibited a directional homologous inhibitory surround (directional-HIS), resulting in an additional inhibition of Imc unit responses when stimuli in the same direction condition. To summarize, the Imc had a directional-HIS that depends on the similarity in the motion direction of the stimuli.

To further analyze the property of the Imc’s directional-HIS, we subtracted the responses of the Imc units in the opposite direction condition from the responses in the same direction condition (SD-OD). This subtraction allowed us to examine the influence of directional-HIS on Imc unit responses at different stimulus distances. The influence of HIS on Imc unit responses at different stimulus distances was illustrated by the blue curves in Fig. 2h and 2k for an example Imc unit and the Imc unit population, respectively. Correlational analysis of these results revealed that the influence of directional-HIS on Imc unit responses gradually diminished with increasing stimulus distance. Furthermore, this effect was evident even when both stimuli were both within the receptive field.

In the above experiments, apart from the influence of directional-HIS on the responses of the Imc units, there existed another inhibitory surround, referred to as “other inhibitory surrounds” (OIS); more precisely, the response of the Imc units in the opposite direction condition is still suppressed compared to when only S1 is present (single). To explore the properties of OIS, we subtracted the response of the Imc unit in the single condition from the response of the Imc unit in the opposite direction condition (OD-Single). The results of the example Imc unit and Imc units population were fitted with the red curves in Fig. 2h and 2k, respectively. The analysis showed that the effect of OIS on the Imc unit gradually increased with increasing stimulus distance and gradually tended to stabilize.

In summary, when the two separate motion stimuli are moving in the same direction, the Imc’s directional-HIS and OIS jointly modulate the response of the Imc units. For directional-HIS, firstly, it depends on the similarity in the motion direction of the stimuli; secondly, the influence on the responses of the Imc unit gradually diminishes with increasing stimulus distance, and this influence occurs when all stimuli are in the Imc unit RF. As the Imc’s directional-HIS is the same as the mutual inhibition in the Itti model and Li’s research (Itti et al., 1998; Li 2016), it can be used to calculate the saliency of the motion direction. For OIS, firstly, it does not depend on the similarity of the motion direction of the stimuli, and secondly, the influence on the responses of Imc units gradually increases until it stabilizes with increasing stimulus distance.

## Modeling and Model validation

### Modeling

The Imc’s directional-HIS depends on the similarity in the motion direction of the stimulus, and this property must be related to the (directly or indirectly) mutual inhibition between homogeneous neurons that can be used to detect a special direction. Existing studies have shown that the only excitatory input to the Imc originates from the Shc of the OT (Li et al., 2007; Schryver & Mysore 2023), and both the OTid and Imc can achieve multimodal perception (visual and auditory) (Schryver & Mysore 2019; Mysore et al., 2010). Previous studies have also found that the OT neurons and Imc neurons don’t exhibit selectivity for the motion direction of targets (Niu et al., 2022; Wang et al., 2022; Wang et al., 2023). Therefore, we propose that the directional-HIS should be computed upstream of the Shc. The retinal afferents of specific retinal ganglion cell types stratify topographically in the OT superficial layers 2–7 except layer 6 (Luksch & Golz 2003), the Shc neurons receive projection from the retinal output layer in the superficial layer of the OT (Wang et al., 2006). The OT 5b layer contains a type of inhibitory neuron (horizontal cells) that receives input from retinal neurons and may constitute local inhibitory circuits within the retino-tectal synapses (Luksch & Golz 2003). In addition to the RGC neurons for motion detection, there are also neurons that may be used to detect other features such as colour, flicker, etc., and the local inhibitory projections within each type of neuron collectively form the Imc’s homologous inhibitory surround (HIS). In summary, we believe that the Imc’s directional-HIS should be computed upstream of the Shc, the calculation of directional-HIS is included here, and it is not limited to directional-HIS. These together constitute the Imc’s HIS and are used to achieve the calculation of stimulus saliency.

The main function of the midbrain network is the stimulus selection. In other words, regardless of the characteristics of the stimulus, only the positions of the most salient stimulus are screened to determine the gaze location at the next moment. The midbrain network achieves stimulus selection depending on inhibitory projections of the Imc neurons to OTid neurons and between the Imc neurons, which are independent of stimulus features and modalities (Schryver & Mysore 2019; Mysore et al., 2010). We believe that this inhibitory projection constitutes the Imc’s non-homologous inhibitory surrounds (non-HIS). Based on the properties of the Imc’s OIS, we consider the Imc’s non-HIS to be the main component of the Imc’s OIS (the Imc’s OIS may contain the Imc’s HIS of other features).

In summary, we believe that the Imc has HIS and non-HIS, in which the former depends on the similarity of stimulus features (e.g., direction of motion, color, etc.), gradually weakens with increasing stimulus distance, and is computed upstream of Shc layer, which can be used for saliency computations of stimuli; in contrast, the latter doesn’t depend on the similarity of stimulus features, gradually enhances with increasing stimulus distance until it stabilizes, and is thought to be generated by inhibitory projections of the Imc to other neurons, which can be used for stimuli selection. Based on the above hypothesis, we constructed a hierarchical coding model of RGC-Shc-Imc, the details were in the model of materials and methods.

### Model validation

In order to ascertain the efficacy of the Imc’s hierarchical neural computing model, a series of tests were conducted, using different visual stimuli. The stimulation video employed in the test consisted of seven frames with a refresh rate of 100 frames/s. The neuron’s membrane potential update time step of the neuron was 1 millisecond, the video size was 100°×100°, and each pixel was 1°×1°. In this model, the diameter of the RF of the model RGC is 1°, the diameter of the RF of the model Shc is 13°, the model Imc only accepts the projection of the model Shc corresponding to its spatial position, and the size of the kernel functions of the projection from the model Imc to model Shc and between the model Imc are 47°×47°. Fig. 3e shows the visual stimulus employed in the model (a translational motion stimulus with a size of 3°×3°), while the responses of the model RGCs are illustrated in Fig. 3f. The responses of the model Shc and Imc populations are presented in Fig. 3g and 3i, respectively. The responses of the model Shc and Imc neurons to motion targets in the four directions are illustrated in Fig. 3h and Fig. 3j, respectively. It can be observed that the model Imc neuron exhibits a stronger response to vertical motion than horizontal motion.

Firstly, the size tuning of the model Imc within Imc’s hierarchical neural computing model was tested. The motion targets of varying sizes were designed through the RF center of the example model Imc, with the mean of the model neuronal responses in the stimulus duration employed to quantify the response of model Imc to motion targets of different sizes. The size tuning curve of the model Imc is illustrated in the red curve in Fig. 4b, while the result of the Imc unit was shown in the black curve in Fig. 4b. The correlation between the two curves is 0.979. This indicates that the model Imc closely matches the results of the size tuning experiment of the Imc unit in the electrophysiology experiment.

**Fig. 4.**
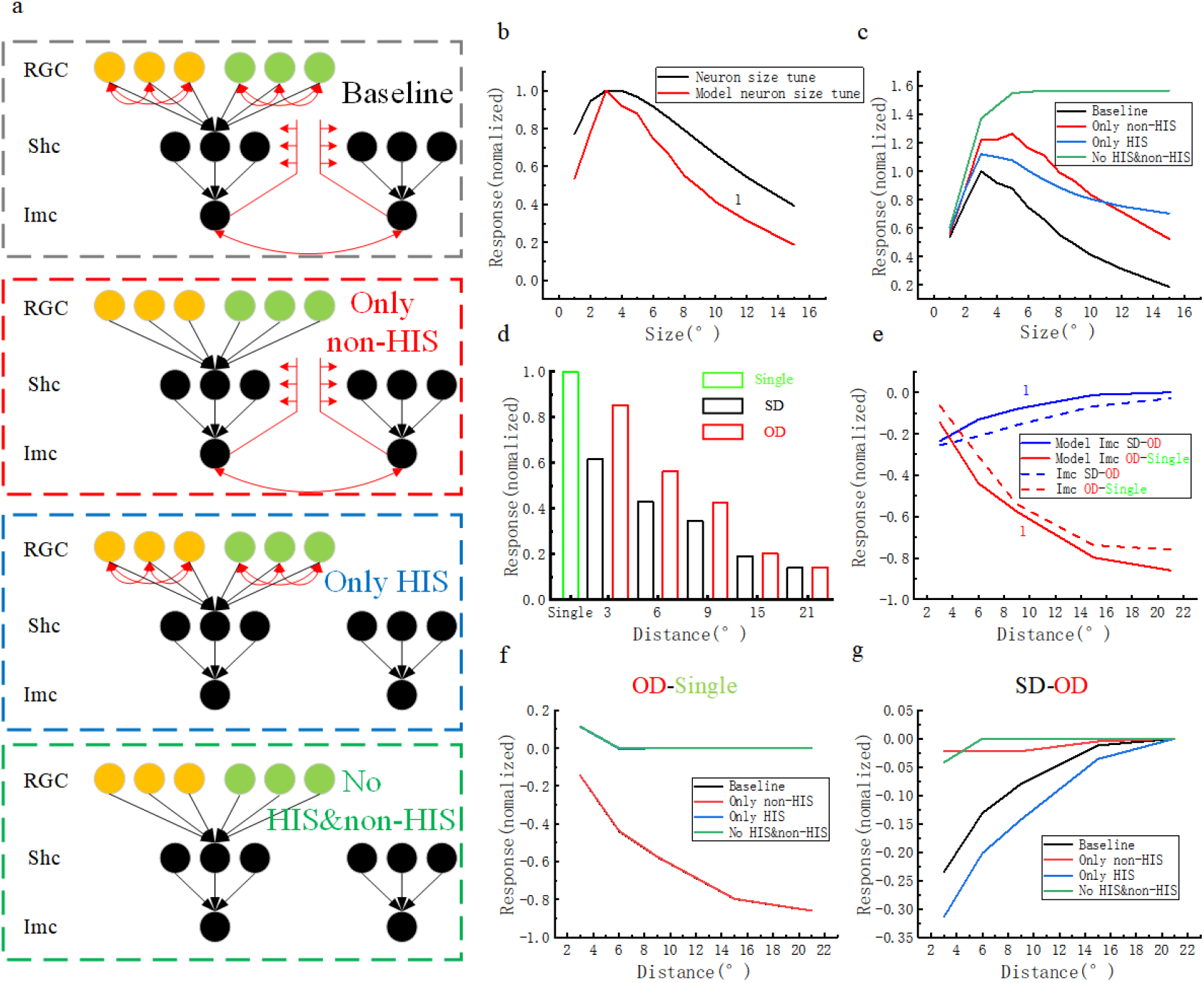
Ablation experiments of hierarchical coding model of RGC-Shc-Imc. a: Schematic diagram of Imc’s hierarchical neural computing model under “Baseline,” “Only non-HIS”, “Only HIS” and “No HIS & non-HIS,” all models built upon a generic “baseline” model. b. The size tuning of the model Imc and Imc units, where the black curve represents the size tuning curve of the Imc unit, the red curve represents the size tuning curve of the model Imc, and the correlation coefficient (Imc unit versus model Imc) is indicated (Spearman test, correlation coefficients is 0.979, p < 0.01, n = 13). c. The size tuning curves of the model Imc after canceling different inhibitory modules (corresponding to Fig. 4a). d. The response of the model Imc to only a single target (green bar), two targets moving in the same direction (black bar), and moving in opposite directions (red bar). e. The influence of Imc’s HIS and non-HIS to Imc unit (dotted line) and model Imc neuron (solid line), where the blue and red curves represent the influence of the Imc’s HIS and non-HIS on the Imc unit and model Imc neuron, respectively; the correlation coefficients (model Imc and Imc unit) are indicated (Spearman test correlation coefficients both are 1, n = 5). f. The response differences of the model Imc between the same direction and the single condition, where black, red, blue, and green curves are the results of the model “baseline”, “only non-HIS”, “only HIS” and “no HIS & non-HIS”, respectively; g: The response differences of the model Imc between same direction and opposite direction condition, where black, red, blue, and green curves are the results of the model “baseline”, “only non-HIS”, “only HIS” and “no HIS & non-HIS”, respectively.

Then, the response of the model Imc to two separate motion stimuli was tested. The stimulus protocols were designed to be identical to those employed in the electrophysiological experiment, as illustrated in Fig. 2a and 2e. The responses of the model Imc were shown in Fig. 4d. The response differences of the model Imc between the same direction and opposite direction conditions were shown by the blue solid curve in Fig. 4e. The correlation of the response differences between the model Imc and Imc unit (blue dotted curve in Fig. 4e) is 1, indicating a strong fit between the model and the electrophysiology experiment results. The response differences of the model Imc between the opposite direction and the single conditions were shown by the red solid curve in Fig. 4e. The correlation of the response differences between the model Imc and Imc unit (red dotted curve in Fig. 4e) is 1, and the model fits the electrophysiology experiment results well.

To further analyze the influence of Imc’s HIS and non-HIS on the model Imc’s response in Fig. 1c, 2a, and 2e stimulation protocols, we conducted ablation experiments on the Imc’s hierarchical neural computational model. Four model configurations were established in the experiment: the complete model (illustrated in the first subplot of Fig. 4a, “Baseline”), blocking the inhibitory projections between homogeneous model RGCs (illustrated in the second subplot of Fig. 4a, “Only non-HIS”), blocking the inhibitory projections of model Imc to other neurons (illustrated in the third subplot of Fig. 4a, “Only HIS”), and blocking all inhibitory projections (illustrated in the fourth subplot of Fig. 4a, “No HIS & non-HIS”).

In the stimulation protocols shown in Fig. 1c, the responses of the Imc units increased and then decreased with increasing target size, and the model Imc also fits this experimental phenomenon (black curve, “Baseline” in Fig. 4c). Compared to “Baseline”, the response of model Imc increased and then decreased with increasing target size. However, the inflection point of its response curve corresponds to a larger target size in “Only non-HIS” (as shown in the red curve in Fig. 4c). For “Only HIS”, the response of the model Imc increased and then decreased with increasing target size, and compared to the “Baseline”, the response of model Imc increased and then decreased with increasing target size, but the attenuation of the response is slower (as shown in the blue curve in Fig. 4c). For “No HIS & non-HIS”, the response of the model Imc increased to a plateau with increasing target size (as shown in the green curve in Fig. 4c). These results indicated that the decrease in Imc response caused by increasing target size was the result of the joint modulation of Imc’s HIS and non-HIS. In the stimulus protocols shown in Fig. 2a and 2e, the response differences of model Imc between the opposite direction condition and the single condition (OD-Single) were as follows: In the case of “Baseline” and “Only non-HIS,” the response of the model Imc was almost equal, and the response becomes more inhibited with increasing stimulus distances. In the case of “Only HIS” and “No HIS & non-HIS,” the responses of the model Imc were also almost equal, and the responses were not inhibited with increasing stimulus distances. The response differences of model Imc between the same direction condition and the opposite direction condition (SD-OD) were as follows: In the case of “Baseline” and “Only HIS,” the additional inhibition described in this study existed in the same direction condition, and the inhibition in “Only HIS” was stronger than that in “Baseline.” The additional inhibition was very weak at “Only non-HIS” and “No HIS & non-HIS”. In summary, Imc’s hierarchical neural computational model constructed in this paper fitted the results of the electrophysiological experiments well, demonstrating that the model has a certain rationality.

The ablation experiments of the hierarchical neural computational model of Imc show that, first, the weakening of the Imc response with increasing target size is the result of the joint modulation of the HIS and non-HIS of Imc; second, this additional inhibition phenomenon, which depended on the similarity of the motion direction of the two stimuli, was mainly caused by the Imc’s HIS, whereas the inhibition phenomenon, which does not depend on the similarity of the motion directions, is caused by the non-HIS of Imc.

## DISCUSSION

In this study, we used Imc neurons as the object of study. First, we found that the response of Imc neurons increased and then decreased with increasing target size by the size-tuned stimulus protocols, and the target size corresponding to the response inflection point was smaller than the range of the ERF of Imc neurons, indicating that there is an inhibitory surround that affects the response of Imc neurons within the range of the ERF of Imc. In further experiments, we used two separate translational motion stimuli as stimulus protocols, subtracted the response of Imc when the two stimulus motions moved in the opposite direction from the response when the motions moved in the same direction (with additional suppression of the Imc response when motions moved in the same direction), and found a HIS of the RF of Imc neurons that showed progressively decreasing suppression of the Imc response with increasing stimulus distance, and the extent of this HIS began within the ERF of Imc. We subtracted the response of Imc when there was only a single stimulus in the centre of the RF from the response when two stimuli moved in opposite directions (the strength of the response to multiple stimuli was less than that to a single stimulus) and found a non-HIS in the RF of the Imc neuron that showed a gradual increase in the inhibitory effect on the Imc response with increasing stimulus distance. The differences between the HIS and the non-HIS of Imc in terms of characteristics are 1. Dependence on similarity of stimulus feature attributes, the former depends on similarity of stimulus feature attributes while the latter does not; 2. Scope of action, the scope of action of the former is local and the effect on the Imc neural response decreases with increasing distance from the centre of the receptive field until it has no effect, while the scope of action of the latter is global and the effect on the Imc neural response increases with increasing distance from the centre of the receptive field until it stabilises. We further analysed the mechanisms of HIS and non-HIS generation in Imc based on existing research on midbrain networks, where the former is calculated by mutual inhibitory projections between similar feature detection neurons upstream of Imc, and the latter by inhibitory projections between Imc neurons. Finally, a hierarchical coding model of RGC-Shc-Imc was constructed, and the plausibility of the HIS and non-HIS generation mechanisms of Imc neurons was verified by correlating the responses of the Imc neuron model with electrophysiological data in different stimulation modes.

### Imc’s HIS is used for saliency computations

The saliency of the stimulus is the basis of stimulus selection for the midbrain network (Knudsen 2018; Schryver & Mysore 2019). It is necessary to determine what the saliency of the stimulus is and where the saliency computation occurs. The saliency of a stimulus is defined as the difference between the stimulus and its surrounds (Yoram 2015; Itti et al., 1998), while the result of the saliency computation for all locations in the visual field is the saliency map. Several existing studies have found a class of neurons in the avian OT (or mammalian SC) that were significantly weaker when the background or surround stimulus and the central stimulus moved in the same direction than when they moved in the opposite direction(Frost et al., 1981;Sun et al., 2002;Zahar et al., 2012;Barchini et al., 2018;Niu et al., 2020;Dutta et al., 2020). Saliency maps of birds in the OT as well as of mammals in the SC (homologous to OT) have also been reported (White et al., 2017). The Itti model, a classical saliency computational model, uses the operation of center-surround differences of homologous features to achieve saliency computation (Itti et al., 1998), that is, the mutual inhibition between homologous neurons. In the OT and V1, mutual inhibition between homogeneous neurons is thought to be a biological mechanism used to calculate saliency (Li 2016). Therefore, combined with the function of the midbrain network used for stimulus selection and the properties of the Imc’s directional-HIS, we believe that the Imc can represent the saliency of the motion direction of the stimulus.

Regarding why the directional-HIS of Imc occurs upstream of Shc, both Otid and Imc are capable of multimodal perception, as the directional-HIS of Imc depends on the similarity of the motion direction of the stimulus. Therefore, Shc are also likely to be multimodal perceptual neurons (Schryver & Mysore 2019; Mysore et al., 2010; Schryver et al., 2020), that is, they can only represent the intensity of the stimulus but not the category of stimulus features. Our previous studies have shown that neurons in the OT are sensitive to translational motion but have no direction selectivity, whereas some RGC neurons have direction selectivity. Previous studies have shown that horizontal cells in the OT5a layer (which receive input from RGCs) are local inhibitory circuits with RGCs and OT neurons (Luksch & Golz 2003), and we consider that Imc’s directional-HIS is calculated by these local inhibitory circuits.

In terms of how neurons achieve the saliency computation of stimuli, we believe that Shc represents the saliency of the stimulus without the inhibitory projection of the Imc, so the saliency computation of the stimulus is divided into two steps: the first step is mutual inhibition between homologous RGCs (the inhibition effect gradually weaken with the increase in stimulus distance), and the second step is a local integration of Shc into the RGC.

### Imc’s non-HIS is used for stimulation selection

The main function of the midbrain network is to achieve stimulus selection, that is, the selection of the gaze location at the next moment. The process of stimulus selection depends on stimulus saliency, and this function is based on the anti-topological inhibition of the Imc to OT and the mutual inhibition between the Imc, which constitutes the Imc’s non-HIS.

### The link between Imc’s inhibitory surrounds and visual receptive field

Some studies have found that the responses of Imc neurons to a bar stimulus of increasing length drop to low values at large lengths (Mahajan & Mysore 2022; Wang & Frost 1991). In further experiments, using extracellular recordings in the Imc coupled with iontophoretic silencing of GABAergic input to the space, the response of OT10 neurons was barely suppressed by longer bar stimulation after OT10 neurons GABA block, whereas the response of the Imc neurons corresponding to that OT10 neurons was still suppressed, although the degree of suppression was reduced compared to that before OT10 neurons GABA block. Based on the current state of research on midbrain network, it is reasonable to assume that the mutually inhibitory projections between Imc neurons are responsible for the suppression of Imc responses in this experiment, as there have been no reports of Imc neurons receiving inhibitory projections from other nuclei. In terms of the stimulus itself, since a longer bar stimulus elicits the response of the Imc neuron at the centre of the bar stimulus, it must elicit the response of other Imc neurons in the vicinity of the bar stimulus, and since inhibitory projections between Imc neurons are thought to be the mechanism by which stimulus selection is achieved at the Imc locus, we can infer that there are inhibitory projections between Imc neurons elicited by a longer bar stimulus. Combined with the conclusions of this paper, it can be concluded that the HIS and non-HIS of the Imc jointly influence the response of Imc neurons in response to longer bar stimuli. The comparison of the responses of the Shc (Fig. 5a) and Imc (Fig. 5b) neuron models before and after the inhibitory projections received by the Shc neuron model in the RGC-Shc-Imc model were switched off further confirms the validity of our conclusions.

**Fig. 5.**
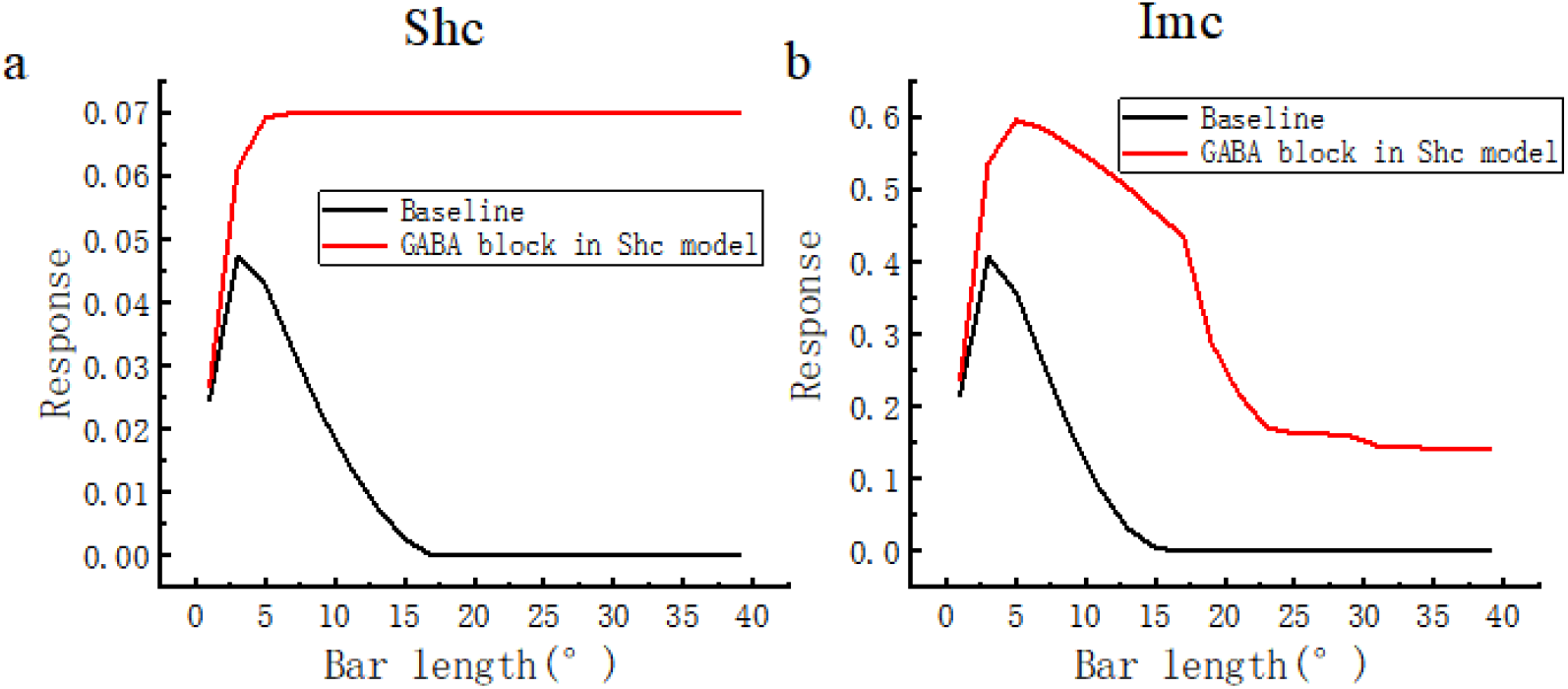
Responses of the model Shc and model Imc before and after GABA block in the model Shc. The horizontal axis in the figure represents the length of the bar stimulus. a. Response of the model Shc before and after GABA block, where the “baseline” (black curve) is the response of the model Shc when the model does not block, and the red curve is the response of the model Shc after GABA block in the model Shc. b. Responses of the model Imc before and after GABA block in model Shc, where the “baseline” (black curve) is the response of the model Imc when the model does not block, and the red curve is the response of model Imc after GABA block in the model Shc.

In the definition of receptive field, the classical receptive field (CRF) is a part of visual space where the presentation or withdrawal of any visual target changes the rate of action potentials in neurons. And in CRF, excitatory receptive field (ERF, or excitatory RF) is the region where presented visual stimuli increase neuron’s spike activities directly, vis-à-vis, inhibitory receptive field (IRF, or inhibitory RF) is the region where presented visual stimuli decrease neuron’s activities (Li et al., 2007). The extraclassical receptive field (eCRF) is defined as the region of visual space where stimuli cannot elicit a spiking response but can modulate the response of a stimulus in the classical receptive field (Henry et al., 2013). The HIS of the receptive fields of Imc neurons in the present study is thought to arise upstream of Imc neurons and is dependent on the similarity of stimulus properties, a feature like the concept of the eCRF. However, the HIS differs from the eCRF in that the firing rate of Imc neurons is enhanced when the target appears within the HIS of Imc alone (the excitatory receptive field of Imc overlaps with the HIS), and thus the HIS of Imc cannot constitute its eCRF. Similarly, Imc’s HIS is even less compatible with the notion of constituting an IRF in Imc’s CRF, because it overlaps with Imc’s eCRF. For the non-HIS of Imc, it mainly functions outside the excitatory receptive field of the Imc neuron and inhibits the response of the Imc neuron when the target is present in this region (Li et al., 2007) and is independent of the target’s attributes, thus the non-HIS of the Imc constitutes the IRF of the Imc.

## Abbreviations

Imc: Isthmi pars magnocellularis
OT: Optic tectum
FR: Firing rate
ERF: Excitatory receptive field
HIS: Homologous inhibitory surround
OIS: Other inhibitory surround
Non-HIS: Non-homologous inhibitory surrounds

## Authors’ contributions

LLQ: Conceptualization, Software, Investigation. CCJ: Writing - original draft. JTW: Software. SWW: Conceptualization, Data curation, Visualization. LS: Validation, Writing - review editing, Supervision. All authors read and approved the final manuscript.

## Availability of data and material

The data that support the findings of this study are available on request from the corresponding author.

## Code availability

The code that supports the findings of this study are also available on request from the corresponding author.

